# Oscillating Hypercapnia Induces Neural Abundant Protein Efflux and Potential Depletion in Health and Chronic Traumatic Brain Injury

**DOI:** 10.64898/2026.04.09.717306

**Authors:** Andrew R. Mayer, Tracey V. Wick, Upasana Nathaniel, Sephira G. Ryman, Divyasree Sasi Kumar, Rebekah Mannix, Samuel Miller, Josef M. Ling, Timothy B. Meier, Kaitlyn Warren, Harm J. van der Horn, Vadim Zotev, Jingshu Wu, Pawani Chauhan

**Affiliations:** The Mind Research Network/Lovelace Biomedical Research Institute, Pete & Nancy Domenici Hall, 1101 Yale Blvd. NE, Albuquerque, NM 87106; Neurology, University of New Mexico School of Medicine, Albuquerque, NM 87131; Psychiatry, University of New Mexico School of Medicine, Albuquerque, NM 87131; Psychology Departments, University of New Mexico School of Medicine, Albuquerque, NM 87131; Boston Children’s Hospital, Boston, MA 02115; Department of Neurosurgery, Medical College of Wisconsin, Milwaukee, WI 53226; Department of Neurology, University of Groningen, University Medical Center Groningen, Groningen, The Netherlands

**Keywords:** Key terms: biomarkers, blood brain barrier, cerebrovasculature, clearance, CSF flow, glymphatic

## Abstract

Emerging preclinical and clinical evidence suggests that low frequency hemodynamic oscillations drive CSF flow, which in turn mediates glymphatic clearance. The current study investigated whether CO_2_-induced low frequency hemodynamic oscillations during magnetic resonance imaging would increase clearance of proteins (glial fibrillary acidic protein, neurofilament light chain, ptau217 and brain-derived tau) from brain to blood, and temporarily improve cognitive performance in individuals with chronic traumatic brain injury (TBI) and age/sex-matched healthy controls. Results indicated that cerebrovascular reactivity, normalized CSF volume, and predicted brain age significantly differed between chronic TBI and controls, while bulk CSF flow differed only at trend levels. Multiple protein concentrations were significantly increased at ∼45 minutes post-hypercapnia, decreased at ∼90 minutes, and returned to pre-hypercapnia levels by ∼150 minutes. Protein efflux was more strongly associated with total CSF volume and total white matter volume rather than cerebrovascular reactivity or bulk CSF flow. Both groups exhibited reduced cognitive interference post-hypercapnia, and hypercapnia associated symptoms quickly returned to baseline levels. In conclusion, hypercapnia temporarily increases clearance of multiple neural abundant proteins into blood, and this effect is moderated by atrophy. Current results suggest that hypercapnia may therapeutically combat pathological protein aggregation post-trauma, and prophylactically during normal aging.

## Introduction

Inefficient clearance of parenchymal waste represents a hallmark of chronic traumatic brain injury (TBI), atypical aging and various other neurodegenerative disorders^1–3^. Metabolic waste, macromolecules and cellular debris are typically cleared through glymphatic/lymphatic pathways (primary) and through the blood-brain barrier (secondary) via transcytosis^4^. The reciprocal relationship between intracranial cerebral blood volume (CBV) and cerebral spinal fluid (CSF) volumes with fixed parenchymal mass was first described over two centuries ago (Monro-Kellie doctrine, Fig. 1a)^5^. More recent preclinical work^6–8^ has emphasized the importance of various vascular factors (Fig. 1b) in the control of CSF flow. CSF flow represents a key and potentially necessary component for glymphatic/lymphatic clearance, as it facilitates subsequent mixing with interstitial solutes, efflux along perivenous, cranial and spinal nerve spaces, and drainage through lymphatic meningeal vessels^4, 6, 9^. Other studies in humans indicate that low frequency hemodynamic oscillations (LFHO) are the primary driver of bulk CSF flow in various brainstem compartments^10–12^, suggesting that exogenous agents which enhance bulk CSF flow can potentially be harnessed for therapeutic purposes^13^.

**Fig. 1:**
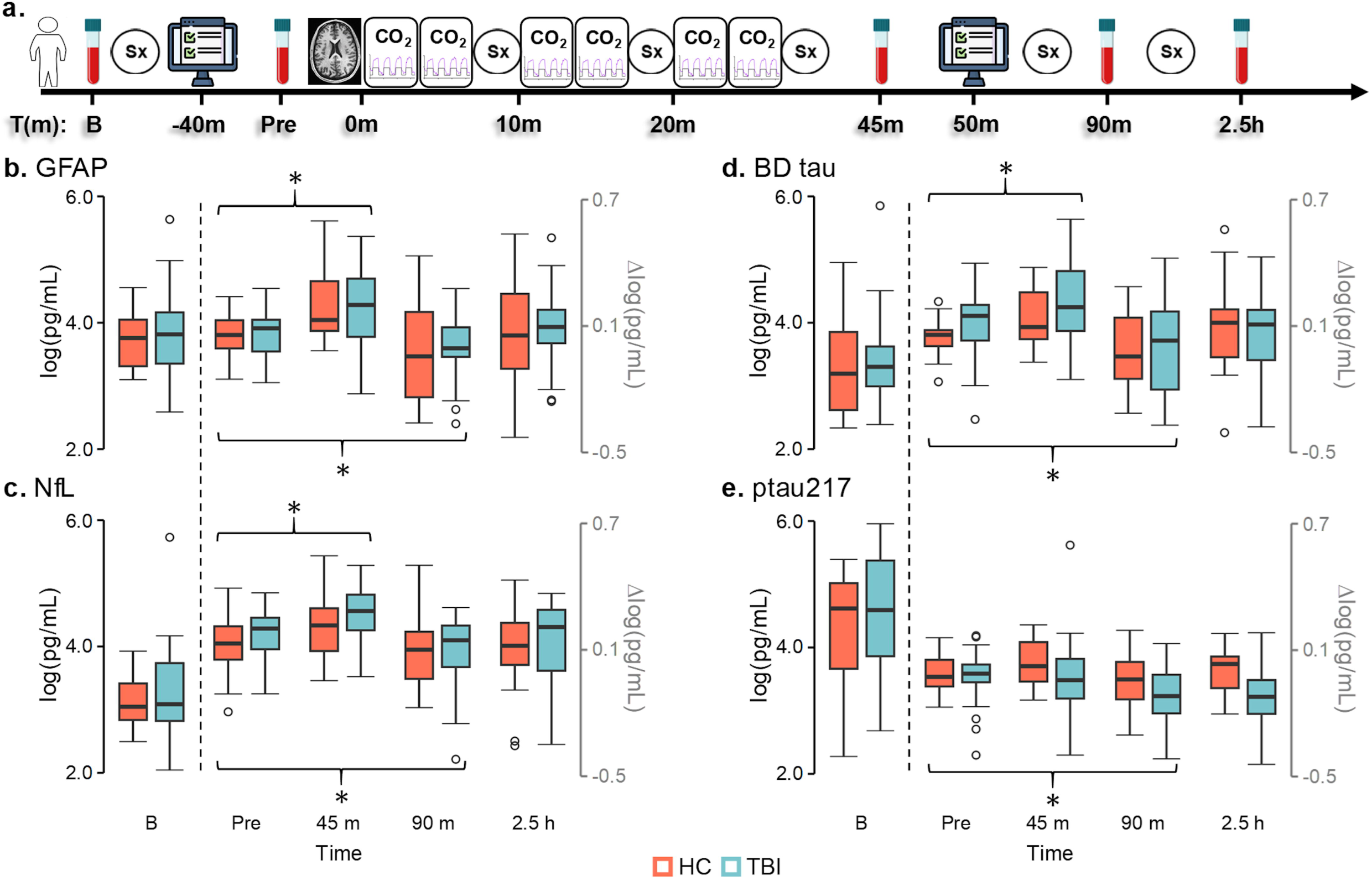
Theoretical and dynamic relationships exist between carbon-dioxide (CO_2_), cerebral blood volume (CBV), cerebral spinal fluid (CSF) flow and protein efflux. CSF occupies ∼13-17% (mustard trapezoid) of intracranial volume, and is inversely proportional (black arrows) to CBV (pink trapezoid; ∼3-7%) assuming constant parenchymal volume (∼80%) and normal brain compliance (Monro-Kellie doctrine; **a**). Global cerebrovascular caliber and tone (**b**) is primarily determined by cerebrovascular reactivity (CVR), vasomotion and physiological (Physio: cardiac and respiratory) fluctuations in order of importance at different frequencies. Typical cerebrovascular caliber range (**b**; top row) is decreased following traumatic brain injury (TBI: **b**; bottom row), as depicted by differences between actual (red) and potential (black) vessel diameter. Dynamic changes in the partial pressure of CO_2_ in arterial blood (measured by end-tidal CO_2_ [ETCO_2_]; **c**) drives CBV changes (ΔCBV; pink arrows) in a non-linear fashion (∼5mmHg ΔETCO_2_ = ∼8% ΔCBV; ∼10mmHG = ∼15% ΔCBV; ∼15 mmHG = ∼25% ΔCBV; maximum ΔCBV∼30%). CBV changes subsequently drive bulk CSF flow (**c**; mustard yellow arrows = ΔCSF) at currently unknown scale (depicted as linear) and maximum value (denoted by X). Changes in CBV (pink arrows)/bulk CSF flow (mustard arrows) are hypothesized to stimulate increased neural abundant protein (NAP) clearance in an unknown dose-dependent fashion (denoted by size of blue arrows; **d**) based on change signal amplitude from the parenchyma (Pa) to blood (Bl). Empirical percent signal change (PSC) data from a single subject demonstrates coupling between gray matter blood-oxygen level dependent response (**e;** pink trace) and bulk CSF flow (**e;** mustard trace on different scale) as a function of ΔETCO_2_.

Waste clearance is greatest during sleep^14, 15^, when large amplitude LFHO occur secondary to global neuronal entrainment^10^ and/or cyclical changes in norepinephrine levels resulting in alterations in vasomotion^6^. In contrast, during wakeful states, changes in CBV are primarily mediated by local neuronal activity within distributed neural networks, with anti-correlated activity occurring across competing networks (e.g., default-mode and executive network)^16^. This diffuse neuronal activity results in small-amplitude changes in global CBV and subsequent small-amplitude changes in vascular-enhanced bulk CSF flow^11, 12^. However, cerebrovascular reactivity (CVR; a primary determinant of cerebrovascular caliber; Fig. 1b) can be parametrically manipulated by the potent vasodilator carbon dioxide (CO_2_) to mimic or exceed the large amplitude changes in CBV that naturally occurs during slow wave sleep (Fig. 1c and 1e). CO_2_ acts directly on smooth muscle cells via H^+^ ions dissolved in CSF from arterial blood (primary mechanism) and secondarily through release of other vasoactive agents^17, 18^. CVR has also been shown to be impaired in both acute and chronic TBI^3, 19–21^ (Fig. 1b). Collectively, these data suggest CO_2_-induced LFHO may be sufficient to enhance waste clearance (Fig. 1d) through advective and diffusion mechanisms associated with increased CSF flow and interstitial solute exchange^13^, and that this process may be impacted by cerebrovascular disease and atrophy following TBI.

The clearance of amyloid proteins from the brain represented a foundational finding establishing the existence of the glymphatic system^22^. Elevated blood protein concentrations have been traditionally equated with protein upregulation and/or cellular breakdown, with each protein linked to individual pathological substrate such as neuronal cell body injury (*ubiquitin* C-terminal hydrolase L1 [UCH-L1]), axonal injury (neurofilament light chain [NfL]), amyloid aggregation ([Aβ] 40 and Aβ42), tauopathies (brain-derived tau, ptau217) or gliosis (glial fibrillary acidic protein [GFAP])^23–26^. Importantly, the concentration gradient for multiple of these proteins has been established to be 9.3 to 130.8% higher in apical tissues and fluids (i.e., CSF and interstitial) relative to blood^27–30^. Several of these neural abundant proteins (NAP) are also increased in the bloodstream following general trauma, anesthetic exposure, cardiac surgery, renal dysfunction and in atypical aging^31–34^. Most recently, blood-based concentration levels for Aβ40, Aβ42, NfL and ptau have been shown to be highest post-sleep^35^, suggesting that NAP may also serve as a surrogate marker for general waste clearance. Collectively these findings suggest that there are multiple, non-exclusive pathways for brain-to-blood efflux, as well as ultimate systemic clearance^36, 37^.

The primary objective of the current study was therefore to investigate whether ∼30 minutes of CO_2_-induced LFHO would be sufficient to promote both NAP efflux and subsequent depletion in patients with chronic TBI (N=22) and healthy controls (HC; N=22). Our a priori hypotheses were that CO_2_-induced LFHO would increase protein efflux, and that this relationship would be moderated by both global tissue volume and the magnitude of CVR. Our second objective was to examine whether ∼30minutes of CO_2_-induced LFHO would be safe and well-tolerated in both healthy individuals and those with chronic TBI. A priori hypotheses were that the majority of participants would be able to safely tolerate the prescribed CO_2_ exposures^18^, but that ∼30 minutes of CO_2_-induced LFHO would not be sufficient to affect performance on complex cognitive control tasks.

## Methods

A prospective cohort design examined effects of hypercapnia on protein efflux (primary endpoints), complex cognition and common symptoms over an approximately 4-hour session (Fig. 2a). Participants with chronic TBI were recruited from the local community and clinics. Age-and sex-matched HC were recruited from the local community based on word of mouth, online databases and posted fliers. Inclusion criteria for the TBI group were 18-68 years old, age-of-injury greater than 18 years, greater than 6 months post-injury, and actively symptomatic. Exclusion criteria included history of neurological, intellectual, or major psychiatric disorders prior to the TBI, active substance abuse/dependence other than cannabis use (determined by urine screen), contraindications for MRI, or non-fluency in English. Exclusion criteria for HC were history of previous TBI with greater than 5-minute loss of consciousness, any neurological diagnosis, psychiatric disorders other than adjustment disorder, autism spectrum disorder, intellectual disability, active substance abuse/dependence, contraindications for MRI or non-fluency in English.

**Fig. 2:**
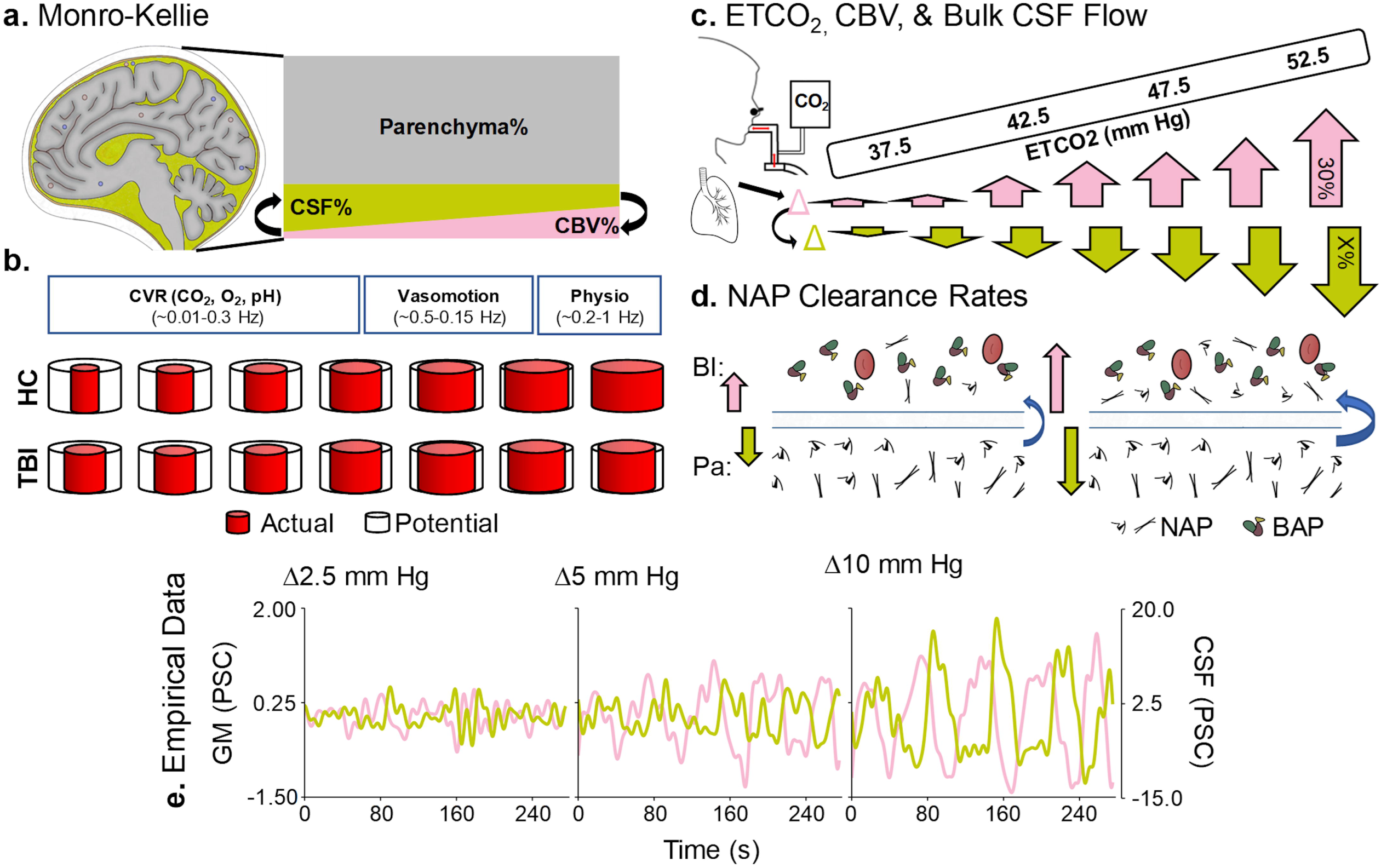
Study design and protein efflux results. Study design (**a**) examining effects of CO_2_ administration on symptoms (Sx), cognitive control (depicted by computer icon) and protein efflux. Carbon-dioxide (CO_2_) challenges were repeatedly administered (first cycle start = Time 0) during blood-oxygen level dependent imaging (depicted with brain icon). Blood samples were acquired at baseline (B), pre-hypercapnia (Pre), 45 minutes (m), 90 minutes and 2.5 hours (h) post-hypercapnia. Glial fibrillary acidic protein (GFAP; **b**), neurofilament light chain (NfL; **c**), brain-derived (BD) tau (**d**), and phosphorylated tau 217 (ptau217; **e**) concentrations (logarithm [log] of picogram per milliliter [pg/mL]) are presented for participants with chronic traumatic brain injury (TBI: light blue) and healthy controls (HC: coral) at baseline in black scale. Change from baseline (Δlog) are depicted in gray scale for all points right of dashed line. Significant post-hypercapnia effects (denoted by brackets with asterisk) associated with both protein efflux (45 minutes; GFAP, NfL and BD tau) and subsequent depletion (90 minutes; all proteins) were observed, followed by return to pre-hypercapnia levels at 2.5 hours (all proteins).

Participants with chronic TBI were classified as experiencing “mild” (N=9) or “moderate-severe” (N=13) injury using a semi-structured interview (Ohio method). However, primary analyses compared all chronic TBI participants to HC in agreement with recent expert panel recommendations^38^.

### Standard Protocol Approvals, Registrations, and Patient Consents

The study was approved by the University of New Mexico Institutional Review Board.

All participants provided informed, written consent.

### Symptom Rating and Cognitive Control Task Description

All participants completed symptom rating (i.e., symptoms of headache, nausea, fogginess and dizziness; 11-point Likert scale) and a cognitive control task (see Supplemental Materials) before and after hypercapnia procedures. Reactive and proactive cognitive control tasks were selected given their critical role for evaluating executive functioning in both health and following TBI^39–41^. Briefly, the task consisted of multisensory (auditory and visual) cues simultaneously presented on either a black (reactive cognitive control task; Fig. 3a) or gray (proactive control task; Fig. 3d) background. A cue prompted participants to attend to either the auditory or visual modality. Cues were followed by simultaneously presented congruent or incongruent multisensory numeric stimuli (targets of “one”, “two” or “three”). During reactive cognitive control, participants pressed a button corresponding to the target number presented in the attended modality while ignoring information in the unattended modality. Cognitive interference was quantified via subtraction of congruent from incongruent trials (i.e., large difference = poor reactive control). During proactive cognitive control (1-back trials), participants were required to hold the attended stimuli in working memory for 1.5 seconds while ignoring the unattended modality, and to only respond to the target upon the subsequent trial. Proactive control was quantified by subtracting congruent 0-back trials from congruent 1-back trials.

**Fig. 3:**
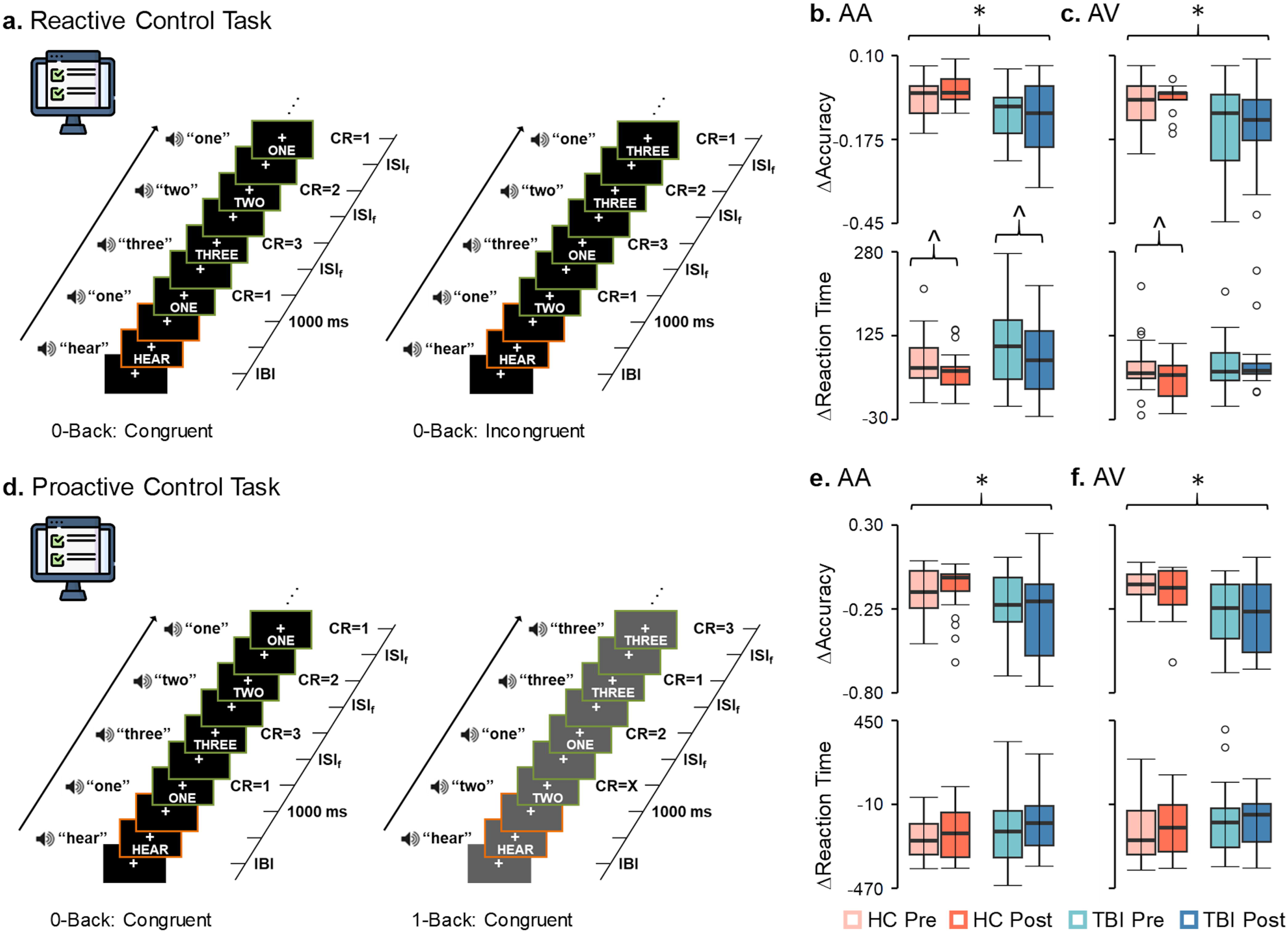
Cognitive control task methods and results. Reactive (**a;** black background) and proactive (**d;** gray background) cognitive control tasks with simultaneously presented auditory and visual numbers (ISI=inter-stimulus interval; IBI=inter-block interval; CR=correct response) were administered pre-and post-hypercapnia to participants with chronic traumatic brain injury (TBI: blue colors) and healthy controls (HC: coral colors). The difference between reaction times (Δ=incongruent 0-back minus congruent 0-back trials) during the reactive cognitive control task significantly decreased indicating less interference) for both groups on attend-auditory trials (AA; **b** bottom row) post-hypercapnia (hypercapnia effects denoted by brackets with carat symbol), and for controls on attend-visual trials (AV; **c** bottom row). In contrast, there were no pre/post-hypercapnia effects during proactive cognitive control (Δ= 1-back congruent minus 0-back congruent trials; **e** and **f** bottom rows). As expected, chronic TBI patients exhibited significantly lower accuracy across both attend-auditory and attend-visual trials relative to healthy controls during both reactive and proactive cognitive control (significant group effects denoted by black brackets with asterisk).

### MRI Hypercapnia Task Description and ETCO_2_ Analyses

All imaging data were acquired on a Siemens 3T scanner with a 32-channel head coil. High resolution T_1_-weighted (voxel size=1.0 mm^3^) images were obtained first followed by 6 runs of single-shot, gradient-echo echoplanar pulse sequence (voxel size=3.0 mm^3^) to quantify bulk CSF flow dynamics during a hypercapnia challenge (Fig. 4a). Each run included 4 separate blocks of increased CO_2_ exposure (35 s duration) interspersed with gas concentrations more typical of atmospheric air (∼30 s duration), for a total of 24 hypercapnia cycles administered across ∼30 minutes. As in previous studies^19^, research staff manually switched between room air (inhalation duration 30±5 s) and the predetermined gas mixture (5% CO_2_, 21% O_2_, balance N_2_) using a Douglas bag, with exhaled CO_2_ data sampled at 1000 Hz. Following pre-processing (see Supplemental Methods), the ETCO_2_ regressor was Min-max normalized to approximate an amplitude of 1 across the entire timeseries, and shifted thirty-five times to capture individual differences in physiology.

**Fig. 4:**
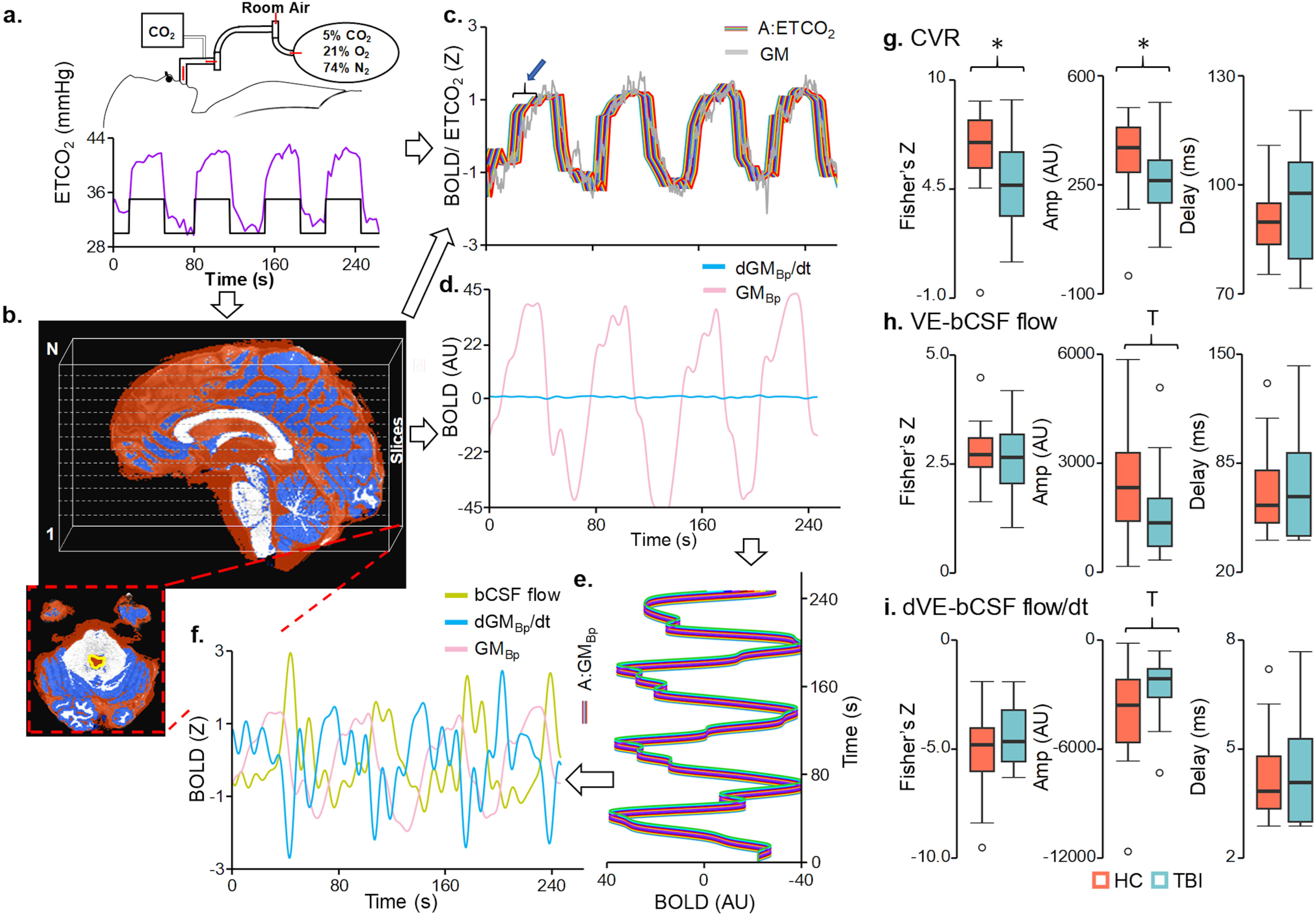
Hypercapnia methods and imaging results. Cerebrovascular reactivity (CVR; **a-c**) and vascular-enhanced bulk cerebral spinal fluid (VE-bCSF; **b, d-f**) flow were quantified using a variety of metrics. Unfiltered global gray matter (GM; gray trace in **c**) as well as bandpass filtered data (GM_Bp_; pink trace in **d**) and its derivative (dGM_Bp_/dt; cyan trace in **d**) were derived from a subject specific mask (dark blue; **b**). The bulk CSF flow signal (bCSF flow; mustard trace in **f**) was derived from a ROI immediately below the 4^th^ ventricle (yellow tracing; **b**). The ETCO_2_, filtered global gray matter signal, and derivative vectors were temporally shifted at various lengths (**c and e**) to account for physiological lags (denoted by blue arrow and bracket in **c**). A series of regressions identified the maximal fit for both cerebrovascular reactivity (GM ROI∼ETCO_2_; **c**) and vascular-enhanced bulk CSF flow (CSF ROI∼GM_Bp_ and CSF ROI∼ dGM_Bp_/dt; **f**) following CO_2_ administration. Participants with chronic traumatic brain injury (TBI: light blue) exhibited a significantly (group effects denoted by asterisk) decreased fit (Fisher’s Z) and amplitude (Amp; arbitrary units [AU]) for CVR metrics (**g**) relative to matched healthy controls (HC: coral). TBI participants exhibited decreased vascular-enhanced bulk CSF flow amplitude (**h;** VE-bCSF flow) at a trend level (denoted by a “T”), with similar results observed for the derivative (**i;** dVE-bCSF flow/dt).

BOLD data during the hypercapnia task were processed using individual CSF and gray matter (GM)-specific pipelines to characterize global CVR^19^ as well as CO_2_-induced bulk CSF flow (Fig. 4b-f; Supplemental Materials and Supplemental Fig. S1)^10, 42^. For CVR quantification, a global GM mask was first created from the T_1_-weighted image (SPM12) using a 60% probability threshold, spatially transformed to the single-band reference image (i.e., native space), and multiplied by the fully preprocessed BOLD data. The array of participant-specific, time-lagged ETCO_2_ vectors was then regressed against the global gray matter signal. The unstandardized beta (i.e., amplitude) from regressor with the maximal fit (i.e., highest absolute Pearson correlation between ETCO_2_ and global GM signal) was selected, followed by a Fisher’s Z-transformation of the Pearson correlation coefficient, and calculation of latency to maximal fit. CSF that has not been exposed to radiofrequency pulses has a higher T2* signal (i.e., flowing into the most inferior slice), while radiofrequency exposed CSF has a lower signal^10^. The CSF-specific voxel-wise preprocessing stream included only time-slice correction, followed by bandpass filtering (0.0008 to 0.10 Hz) tailored to CVR cycle length, detrending and truncation to remove edge artifacts. A CSF region of interest within the cervical central canal inferior to the medulla oblongata was identified via a semi-automated process. To assess CO_2_-induced bulk CSF flow, the fully preprocessed voxel-wise data were similarly bandpass filtered, detrended and truncated, and multiplied by the subject specific GM mask to form a single vector. The bandpassed GM vector was time-shifted 83 times to form an array of regressors (0 to 40.32 s). The single vector was also numerically differentiated to create a GM derivative to specifically quantify the relationship between changes in CBV and bulk CSF flow, as well as for quantifiable quality assurance metrics (see Supplemental Methods and Supplemental Fig. S1). The derivative was then time-shifted 6 times to create a regressor array spanning from 0 to 3.36s. Both the bandpassed GM and derivative arrays were subsequently regressed on the CSF flow vector to quantify CO_2_-induced bulk CSF flow using identical metrics as the CVR analyses (i.e., normalized amplitude, Fisher’s Z and latency of maximal fit).

Total GM, WM and CSF volumes were obtained from the T_1_-weighted image (FreeSurfer pipeline) and normalized by total intracranial volume as primary structural metrics. Brain age (Supplemental Fig. S2) was estimated using a deep learning network (DeepBrainNet) architecture^43^. The predicted brain age difference (PAD) was calculated by subtracting actual subject age from the median value of all slice-wise predictions, and subsequently served as a secondary measure of overall brain structure health.

### Blood Processing

Blood samples were collected at baseline, pre-hypercapnia, 45 minutes, 90 minutes and 2.5 hours post-hypercapnia (Fig. 2a). Protein concentrations in plasma samples were measured in duplicate with a Simoa HD-X Analyzer using the NP4D kit (GFAP, NfL, UCH-L1 and brain-derived tau) and ALZpath ptau217 assay. All blood-based biomarker data were natural log-transformed prior to analyses.

### Statistical Analysis

Chi-square tests or Generalized Linear Models (GLM) were used to evaluate group differences in demographics using IBM SPSS Statistics package. Our primary analyses used protein change scores from baseline (4 change measurements) and Group×Time Linear Mixed Effects (LME) models to examine for effects of hypercapnia on protein efflux (i.e., main effect of Time). Sample size and power for the current study were determined based on iterative testing of effects associated with protein efflux during data collection (main effect of Time) rather than differences between TBI and HC groups. The safety/tolerability of the hypercapnia protocol was similarly evaluated using change scores (symptoms; 5 measurements) and a Group×Time LME, whereas pre/post changes in reactive or proactive cognitive control (pre/post reaction times and accuracy) were examined with Group×Time LME models. Auditory and visual trials were analyzed separately given previously reported differences in performance based on modality for focused attention^44^. LME models were also used to examine for group differences in CVR or bulk CSF flow metrics, as well as normalized structural volumes (primary) and predicted brain age (secondary). Finally, separate multiple regression with partial correlations assessed for relationships between CVR and bulk CSF flow metrics (fit and amplitude) with protein efflux, or between normalized structural volumes and protein efflux.

Sensitivity analyses were conducted to ensure that analyses with or without outliers did not change statistical conclusions for primary findings (e.g., significant to non-significant, trend to non-significant, or non-significant to trend or significant). Any deviations noted during sensitivity analyses are reported in the results section. Planned primary analyses for protein efflux were corrected for family-wise error using the Bonferroni method. Interactions were examined with simple effects testing.

## Results

### Demographics and Symptom Characterization

A total of 22 individuals with chronic TBI (9 “mild” and 13 “moderate-severe”) and 22 HC completed all hypercapnia exposures (24 cycles across 6 MRI runs) for a completion rate of 95.7%. Two participants (4.3% of sample) were unable to tolerate the hypercapnia equipment (mask inside of scanner) due to claustrophobia rather than the CO_2_ exposure itself. Participants who completed the hypercapnia task did not differ on biological sex or age between the TBI and HC groups (all *p*’s >0.05; Table 1). The total CO_2_ exposure summed across the 6 individual runs did not significantly differ as a function of Group (*p*=0.124; Cohen’s d=-0.47), sex or age.

**Table 1:** Participant demographics and symptom summary scores (i.e., Likert ratings of headache, nausea, fogginess and dizziness) across the entire hypercapnia challenge.

|  | TBI (N=22) | HC (N=22) | *p*-value |
| --- | --- | --- | --- |
| Age | 42.41±14.44 | 42.73±11.26 | 0.934 |
| Sex (%Male) | 14(63.6%) | 13(59.1%) | 0.757 |
| **Injury Characteristics** |  |  |  |
| Severity (“mTBI”/ “msTBI”) | 9(40.9%)/13(59.1%) | NA |  |
| YPI (years post-injury) | 3.7(1.6-6.1) | NA |  |
| **Symptoms** |  |  |  |
| B | 0(0-3.25) | 0(0-1) | <0.001 |
| ΔCO<sub>2</sub>2 | 0(-1.25-3) | 0(-0.25-0) | 0.884 |
| ΔCO<sub>2</sub>4 | 0(-1.25-2.25) | 0(-0.25-0) | 0.432 |
| ΔCO<sub>2</sub>6 | 1(0-4) | 0(-0.25-0.25) | 0.052 |
| ΔBl<sub>4</sub> | 0(-1-0) | 0(-0.25-0) | 0.851 |
| ΔBl<sub>5</sub> | 0(-2-0) | 0(-1-0) | 0.170 |

Notes: “mTBI”= “mild” TBI; “msTBI”= “moderate-to-severe” TBI; NA=not applicable; B=baseline; Δ=change score between the timepoint provided and baseline; ΔCO<sub>2</sub>2=symptom rating after second block of hypercapnia challenge; ΔCO<sub>2</sub>4=symptom rating after fourth block of hypercapnia challenge; ΔCO<sub>2</sub>6=symptom rating after sixth and final block of hypercapnia challenge; ΔBl<sub>4</sub>=symptom rating after fourth blood draw; ΔBl<sub>5</sub>=symptom rating after fifth and final blood draw. All data are reported in median and IQR (denoted by parentheses) or mean plus standard deviation (denoted by ±).

### Effects of Hypercapnia on Protein Efflux

Blood collection was successful for 98.2% of attempts across the 44 individuals who completed the imaging protocol (see Supplemental Results). Quality assurance protocols yielded low failure rates for GFAP (2.9%), NfL (3.9%), brain-derived tau (1.5%) and ptau217 (2.9%), whereas the majority of UCH-L1 samples (56.4%) exhibited poor data quality as denoted by high coefficients of variation (51.5%) or samples that were below levels of detection (4.9%). As a result, UCH-L1 data were not analyzed further. Trend group differences (TBI>HC) were observed for baseline GFAP (F=3.32; *p*=0.077; Fig. 2b), NfL (F=3.71; *p*=0.062; Fig. 2c) and brain-derived tau (F=3.15; *p*=0.085; Fig. 2d), though sensitivity analyses indicated that baseline differences in GFAP and NfL were strongly influenced by a single individual with chronic TBI (see Supplemental Results for effect size calculations).

Change scores from baseline were calculated for all 4 proteins. A series of 2×4 (Group [HC vs TBI] × Time [pre-hypercapnia, 45 minutes post-hypercapnia, 90 minutes post-hypercapnia, 2.5 hours post-hypercapnia]) LME analyses indicated significant main effect of Time for GFAP, NfL, brain-derived tau and ptau217 (all Bonferroni corrected *p’s*<0.0125) change scores. Protein efflux occurred at 45 minutes post-hypercapnia for GFAP (*p*<0.001), NfL (*p*=0.002) and brain-derived tau (*p*=0.001) relative to the pre-hypercapnia timepoint, whereas ptau217 efflux was not significant (*p*=0.127). Significant evidence of decreased GFAP (*p*=0.011), NfL (*p*=0.021), brain-derived tau (*p*=0.002) and ptau217 (*p*=0.028) was observed at 90 minutes post-hypercapnia relative to pre-hypercapnia (Fig. 2b-e) levels, with all proteins returning to pre-hypercapnia levels by 2.5 hours (all *p*’s>0.11). The main effect of Group and Group×Time interactions for all 4 proteins were non-significant (all *p*’s>0.10). Effect size ranges for differential protein efflux (d=-0.39 to-0.43) and depletion (d=-0.15 to-0.48) as a function of Group (chronic TBI>HC) were all in the small to medium range.

Supplemental Table 1 presents means and standard deviations for baseline protein levels, as well as change scores for HC, all chronic TBI participants, chronic “mild” TBI participants and chronic “moderate-to-severe” TBI participants.

### Safety and Tolerability of the Hypercapnia Protocol

Significant (*^2^*=12.38, *p*<0.001) increases in baseline symptoms (headache, nausea, fogginess and dizziness; Supplemental Fig. S3; Table 1) existed for TBI relative to HC. A 2×5 (Group [HC vs TBI] × Time [post-second hypercapnia block, post-fourth hypercapnia block, post-sixth hypercapnia block, post-fourth blood draw, post-sixth blood draw]) LME model using symptom change scores from baseline indicated significant main effect of Time (F=4.53, *p*=0.002) and a significant Group×Time interaction (F=4.44, *p*=0.002). However, only trend differences existed between TBI and HC participants following the sixth and last hypercapnia run (F=4.01, *p=*0.052; all other *p*’s>0.10). Symptom ratings peaked following the 6^th^ relative to 2^nd^ hypercapnia run for both groups (*p*=0.014), and were also significantly different relative to all other post-hypercapnia assessments (all *p*’s<0.007).

### Multisensory Proactive and Reactive Cognitive Control Task Effects

A series of 2×2 (Group [HC vs TBI] × Time [Pre-CVR Task vs Post-CVR Task]) LME models examined for effects of extended hypercapnia on reactive (accuracy and reaction time; Fig. 3b-c) and proactive (Fig. 3e-f) cognitive control. Accuracy was decreased for individuals with TBI relative to HC during reactive cognitive control for both attend-auditory (F=9.874; *p*=0.003) and attend-visual (F=7.276; *p*=0.010) conditions, with no other main effects or interactions (all *p*’s>0.05). The main effect of Time was significant for the attend-auditory condition (F=4.542; *p*=0.039), with significantly improved reactive cognitive control post-hypercapnia task across both TBI and HC participants. A significant Group×Time interaction (F=6.232; *p*=0.017) was present within the attend-visual condition, with significantly faster reactive cognitive control post-hypercapnia for HC (F=5.765; *p*=0.026) and null effects for chronic TBI (*p*>0.05).

During proactive control, significant main effects of Group (HC> chronic TBI) existed for accuracy with both attend-auditory (F=4.159; *p*=0.048) and attend-visual (F=9.653; *p*=0.003) conditions, with non-significant Time or Group×Time interactions (all *p’*s>0.05). No significant main effects or interactions were present for reaction time data during proactive cognitive control (all *p*’s>0.05).

### CVR and Bulk CSF Flow Coupling Results

The majority (90.5%) of the individual CVR runs exhibited significant evidence of anti-correlation (>0.025% of null distribution) between the derivative of the global GM signal and bulk CSF flow (see Supplemental Results). The sum of framewise displacement across all runs was higher for TBI relative to controls at a trend level using an LME model with Group as the single factor (F=3.25; *p*=0.079). LME models with group, sex, age and mean framewise displacement as covariates therefore examined group differences in CVR and CO_2_-induced bulk CSF flow coupling summed across all 6 runs to assess for cumulative effects of the hypercapnia protocol.

Results from these LME models indicated significantly greater CVR (Fig. 4g) maximal fit (F=4.69; *p*=0.036; Cohen’s d=0.66) and normalized amplitude (F=4.45; *p*=0.041; Cohen’s d=0.64) for HC relative to TBI, whereas CVR latency was not significant (*p*=0.953; Cohen’s d=-0.02). In contrast, no significant Group effects existed for CO_2_-induced bulk CSF flow fit or latency for either the bandpass filtered data (Fig. 4h) or its derivative (Fig. 4i). Specifically, CO_2_-induced bulk CSF flow amplitude was different at trend level only for the Group effect (F=3.86; *p*=0.057; HC>TBI; Cohen’s d=0.61), whereas small effect sizes were observed for latency (Cohen’s d=0.14) and fit (Cohen’s d=0.09). Similarly, amplitude was different at trend level (F=3.86; *p*=0.057; HC<TBI; Cohen’s d=-0.61) for the derivative (more negative for HC), with small effect sizes for latency (Cohen’s d=-0.20) and fit (Cohen’s d=-0.34). Age was significantly and positively related to both summed derivative fit (F=8.76; r=0.44; *p*=0.005) and normalized derivative amplitude (F=5.08; r=0.37; *p*=0.030).

### Structural Imaging Results

A series of LME models comparing tissue volumes (Fig. 5a-c) across Group with sex and age as additional covariates indicated that normalized CSF volume was significantly increased for participants with chronic TBI relative to HC (F=7.31; *p*=0.010; Cohen’s d=-0.81). In contrast, normalized WM (*p*=0.180; Cohen’s d=0.41) and GM (*p*=0.169; Cohen’s d=0.42) volumes did not show a significant main effect of Group at current sample sizes (*p*>0.10), although medium-to-large effect sizes were present. A similar LME model indicated that predicted brain age (Supplemental Fig. S2) was significantly higher for participants with chronic TBI relative to HC (F=4.84; *p*=0.034; Cohen’s d=-0.65).

**Fig. 5:**
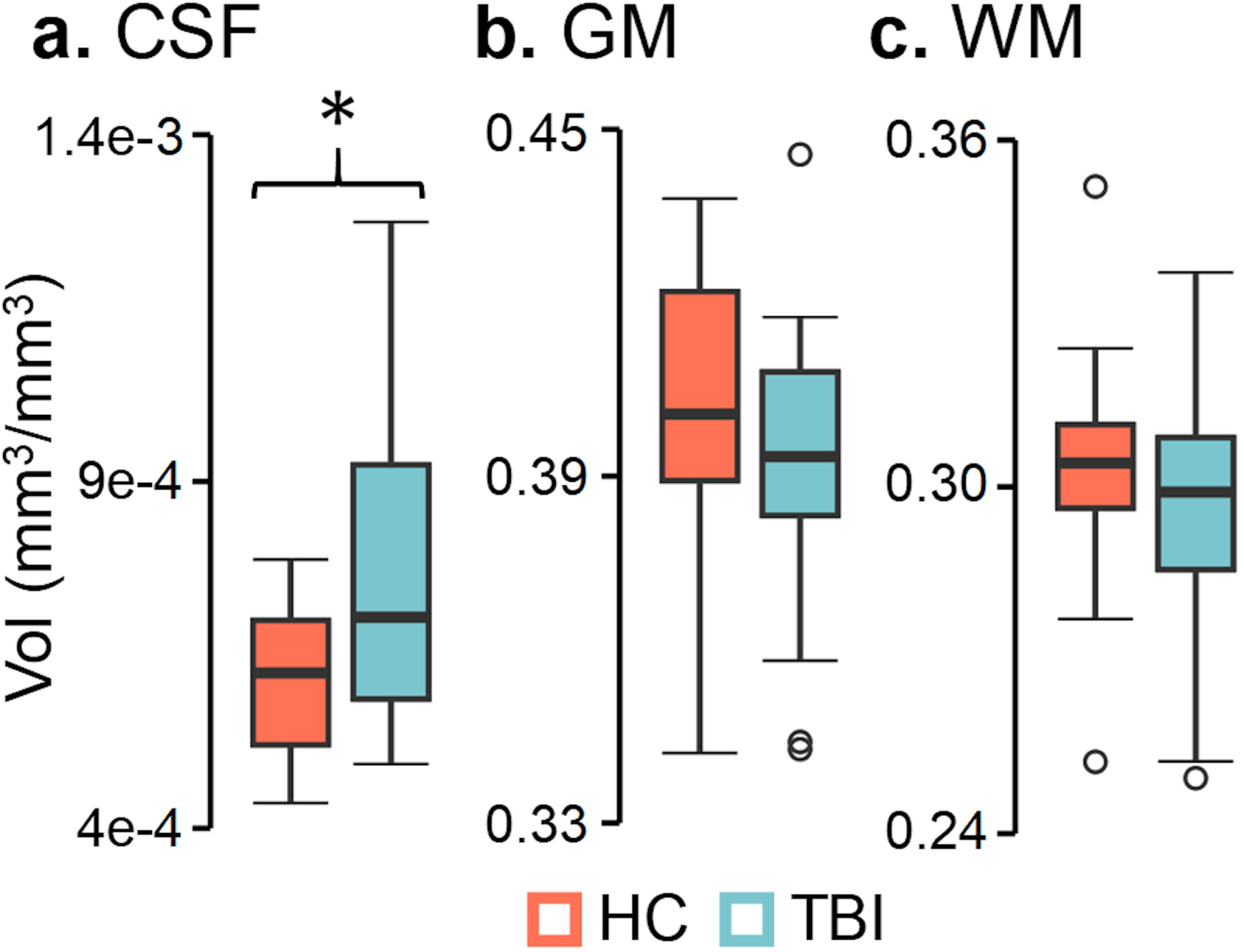
Structural imaging results. Participants with chronic traumatic brain injury (TBI: light blue) exhibited significantly (group effects denoted by asterisk) increased normalized cerebral spinal fluid (CSF; **a**) volume (Vol) relative to healthy controls (HC; coral), whereas normalized gray (GM; **b**) and white (WM; **c**) volumes were not significantly different.

### Relationships between Imaging Results and Protein Efflux

Separate multiple regression analyses investigated associations between fit/amplitude from CVR and CO_2_-induced bulk CSF flow (independent variables) with protein efflux (change scores 45 minutes post-hypercapnia), as well as between structural volume metrics (independent variables: normalized CSF, GM and WM) and protein efflux. Omnibus results from the vascular/bulk CSF flow metric models were significant only for ptau217 (Fig. 6a; *p*=0.010; R^2^ change=30.9%), with significant positive partial correlations observed between ptau efflux and CVR fit (ρXY⋅Z=0.34) and CO_2_-induced bulk CSF flow amplitude (ρXY⋅Z=0.35).

**Fig. 6:**
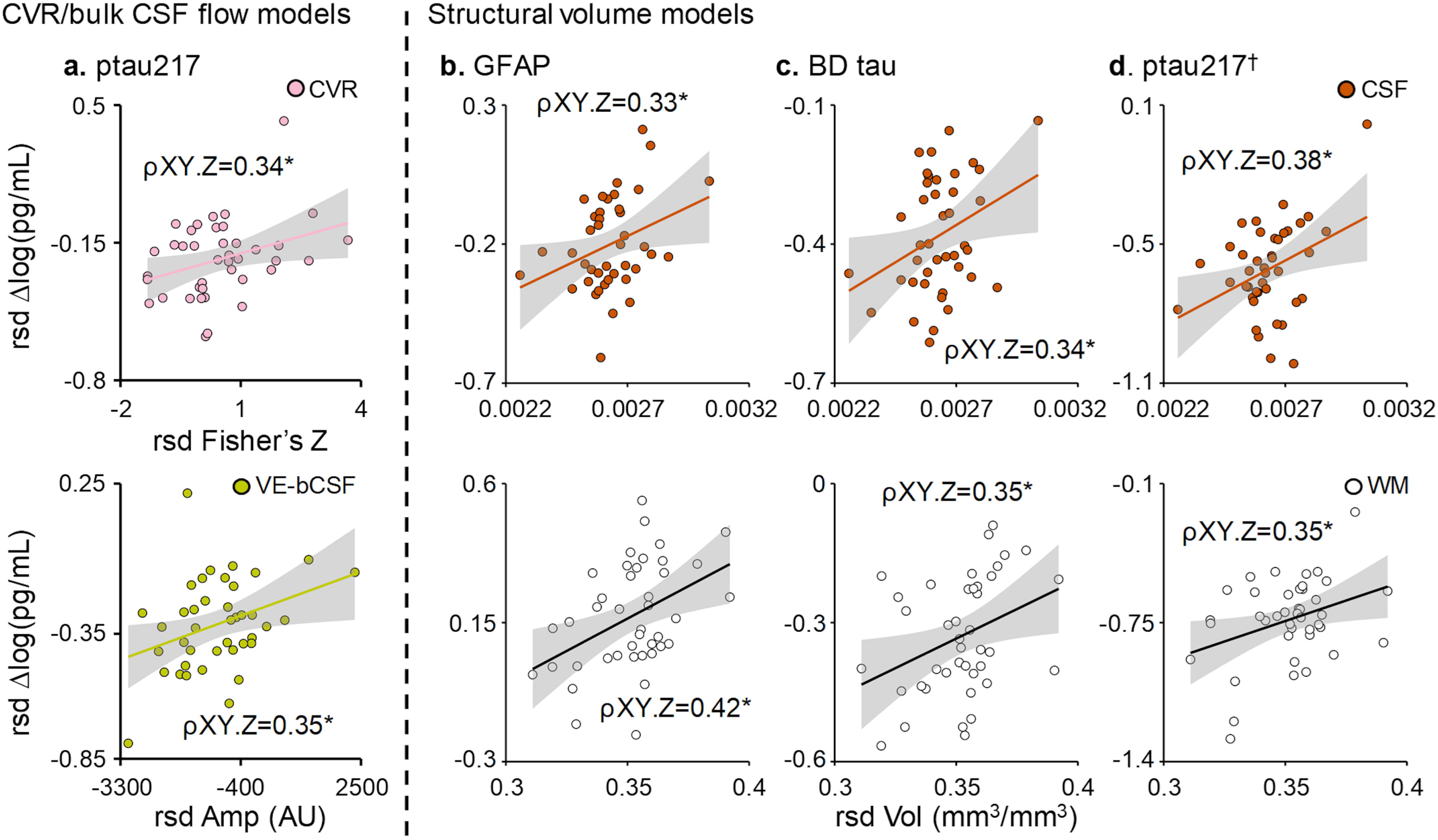
Relationship between imaging metrics and protein efflux. Significant and unique associations (denoted by partial correlation = ρXY.Z; all data graphed in residualized [rsd] form respective to all other variables) were observed between cerebrovascular reactivity (CVR) fit (Fisher’s Z; pink circles) and phosphorylated tau 217 (ptau217; **a**) efflux (logarithm [log] of picogram per milliliter [pg/mL]) at 45 minutes post-hypercapnia, as well as between CO_2_-induced bulk cerebral spinal fluid (CSF) amplitude (Amp; arbitrary units [AU]; mustard circles) with ptau217. Associations between structural volumes and protein efflux were significant for glial fibrillary acidic protein (GFAP; **b**) and brain-derived (BD) tau (**c**), with both CSF volume (orange circles) and white matter volume (white circles) each uniquely associated with protein efflux. Similar structural relationships existed for ptau217 (**d**), although the overall model was not significant (denoted by ^†^).

In contrast, omnibus results for tissue volume models (Fig. 6b-d) were significant for GFAP (*p*=0.024; R^2^ change=26.3%) and brain-derived tau (*p*=0.039; R^2^ change=23.9%) efflux, with significant positive partial correlations observed between normalized CSF (GFAP ρXY⋅Z=0.33; brain-derived tau ρXY⋅Z=0.34) and white matter (GFAP ρXY⋅Z=0.42; brain-derived tau ρXY⋅Z=0.35) volumes with the magnitude of protein efflux. Although the overall model for ptau217 was not significant (*p*>0.05), a similar magnitude of partial correlations were observed for CSF (ρXY⋅Z=0.38) and white matter (ρXY⋅Z=0.35) volumes.

## Discussion

The dynamic, inverse relationship that exists between global CBV and CSF volumes within the fixed cranial vault was originally posited over two hundred years ago^5^, but the importance of vascular-enhanced CSF flow in glymphatic functioning has only recently emerged^6–8^. Consistent with a priori hypotheses, CO_2_-induced LFHO were effective for increasing the efflux of neural abundant protein concentrations in the blood, which may represent a surrogate marker of brain waste clearance^36^. This temporary increase was subsequently followed by a reduction in blood protein concentrations below pre-hypercapnia levels, providing preliminary evidence of a putative reduction in parenchymal protein concentrations following ∼30 minutes of oscillating CO_2_ exposure. Current imaging results indicated that individuals with chronic TBI exhibited reduced CVR and increased atrophy relative to matched controls, but no group differences existed for CO_2_-induced bulk CSF flow at our limited sample sizes. The magnitude of protein efflux was more strongly associated with CSF and white matter tissue volumes, although ptau217 was also associated with both vascular functioning and bulk CSF flow coupling. Collectively, current and previous^45^ results suggest that CO_2_-induced LFHO enhance waste clearance across multiple neurodegenerative conditions, and may therefore have therapeutic indications.

In normal tissue, healthy epithelia (e.g., blood-brain and blood-CSF barriers) typically restrict the exchange of proteins from apical and basolateral spaces through continuous non-fenestrated endothelial cells and tight junctions, with protein exchange occurring through highly regulated transcytosis^46, 47^. The glymphatic/lymphatic systems therefore represent the primary pathway for clearance of waste products into the bloodstream through both advective and diffusion processes^4, 6, 9^. Glymphatic/lymphatic clearance is dependent on both macroscopic and microscopic mechanisms which vary across gray and white matter tissue, but CSF influx is recognized as a necessary component as it facilitates mixing of interstitial solutes and subsequent waste removal. Human^10–12^ and rodent^6–8^ data suggest that LFHO respectively govern both global (e.g., laminar flow in the 4^th^ ventricle) and local (e.g., at the level of neurovascular unit) CSF flow dynamics. Thus, increases in both global (e.g., cerebrovascular reactivity) and local (e.g., following neurovascular coupling) cerebral blood flow/volume not only provide needed metabolic substrates, but also play a critical role in indirectly promoting waste clearance via increased CSF flow^7, 8^.

Prescribed CO_2_ can be used to mimic the oscillating levels of global parenchymal engorgement that occur naturally during slow wave sleep secondary to neuronal entrainment^10^ or vasomotion^6^, and have been shown to drive glymphatic clearance in rodent studies. Importantly, the majority (90.9%) of individual hypercapnia sessions (i.e. 6 runs per individual; 264 runs in total) resulted in strong anti-correlation between the derivative of CO_2_-induced LFHO and bulk CSF flow, indicative of the fundamental inverse relationship (i.e., increased blood volume=decreased CSF volume) that exists for global fluid dynamics within the fixed cranial vault^5^. The tight coupling that occurs between CO_2_-induced LFHO and bulk CSF flow did not differ between HC and individuals with chronic TBI, although differences in vascular-induced CSF flow amplitude changes existed at trend level. Consistent with a priori hypotheses, GFAP, NfL and brain derived tau concentrations were increased in the blood stream ∼45 minutes after the hypercapnia challenge across both groups of participants, followed by evidence of potential parenchymal depletion (i.e., reduced plasma protein levels) at ∼90 minutes post-hypercapnia for all measured proteins. The kinetics of protein efflux and depletion were temporally uniform regardless of protein molecular weight at the current temporal sampling regimen, a finding more commonly associated with advection during glymphatic clearance^4^. Although UCH-L1 was quantified, results indicated poor data quality, similar to previous studies with this assay for “mild” TBI in more acute settings^48, 49^.

TBI alters endothelial and smooth muscle cell function, changes capillary distensibility, reduces capillary recruitment and cerebral blood flow, and causes pericyte loss^3, 50, 51^. Post-traumatic vascular pathology has been histologically confirmed even after controlling for vascular risk factors and vascular disease burden^52^. The traumatically-compromised cerebrovasculature has been linked to reduced waste clearance^1, 3^, which further disrupts the neurovascular unit through positive feedback loops (“double-hit” hypothesis) and increases the likelihood of prion-like spreading. Hypoperfusion has traditionally been proposed as the primary link between cerebrovascular dysfunction and reduced waste clearance, but previous^3, 20, 21^ and current findings suggest CVR is also reduced post-TBI, and therefore represents a second potential mechanism. Specifically, a limited vessel caliber range may result in delayed or reduced cerebral blood flow, as well as restrict the magnitude of subsequent bulk CSF flow (Fig. 1b-c), both of which **may** negatively impact waste clearance in the long-term. However, contrary to our a priori predictions, there was only a significant relationship between CVR and vascular-induced bulk CSF flow with ptau217 rather than all proteins.

Multiple studies^53–56^ have indicated progressive tissue loss (i.e., neurodegeneration) following TBI, which is greater in white (+5.97 years) relative to gray (+4.66 years) matter and is associated with blood protein concentration levels. CSF volumes and predicted brain age were significantly increased in the chronic TBI cohort relative to the control sample, and global CSF and white matter volumes accounted for unique variance for GFAP, brain-derived tau and ptau217 efflux. These findings preliminarily suggest that CO_2_-induced protein efflux may be more dependent on global parenchymal tissue to fluid ratios (Monro-Kellie doctrine) rather than fluid dynamics at the neurovascular unit level (i.e., local interactions between changes in vessel caliber and CSF flow)^6–8^. However, these findings require replication in larger sample sizes and other neurodegenerative conditions. Moreover, current results are likely only pertinent to acute alterations in fluid dynamics resulting from CO_2_ administration rather than the long-term relationships that exist between cerebrovascular dysfunction and abnormal protein aggregation^1,3^. Therefore, future preclinical and clinical studies are required to examine the role of these individual biological mechanisms on waste clearance in more detail.

For example, systemic clearance through renal and hepatic pathways represents a frequently overlooked factor for understanding protein concentration levels in blood^36^. Clearance via renal filtration occurs exponentially faster for proteins with lower molecular weights when other molecular factors (charge and size) are held constant, whereas >60 kDA proteins are typically catabolized hepatically^57, 58^. Moreover, although CO_2_ is a potent vasodilator in most tissue beds, it acts as a vasoconstrictor in the kidneys and results in both decreased blood flow and increased tissue resistance in healthy individuals^59, 60^. Similarly, recent evidence suggests that there may be at least two separate pathways for glymphatic clearance from parenchymal tissue versus the CSF that are at least partially dependent on molecular weight, with larger proteins potentially captured and cleared through lymph nodes^37^. Thus, future preclinical studies are also required to determine how vascular-enhanced manipulations of bulk CSF flow affect different glymphatic clearance pathways and compounds of various molecular weights.

Previous preclinical studies have demonstrated that glymphatic impairment decreases the release of waste products into the blood stream during the acute phases of TBI, theoretically increasing cellular exposure to these toxins^61^. Hypercapnia may therefore represent a generalized therapy that can be prescribed to enhance waste clearance early post-injury, when treatments are most likely to be more effective. Similarly, the therapeutic administration of hypercapnia could also reduce post-traumatic toxic protein concentrations in the parenchyma during the chronic injury phase, as such proteins can remain elevated months-to-years post-injury and are associated with increased atrophy^53–55^. Finally, regularly prescribed hypercapnia sessions may also augment homeostatic waste clearance given that sleep is frequently disrupted following TBI or in atypical aging^62^.

Although individuals with TBI and HC experienced a temporary increase in symptoms during the hypercapnia session, total symptom burden quickly returned to baseline levels, suggesting that the hypercapnia procedures were safe and well tolerated. No other adverse events were observed during the study. Contrary to a priori hypotheses of a null effect, performance on a reactive cognitive control task improved after approximately 30 minutes of oscillating hypercapnia, with evidence of reduced interference from conflicting stimuli in both individuals with chronic TBI and HC. In contrast, there were no significant post-hypercapnia changes in proactive cognitive control for either group. Individuals with TBI exhibited the expected impairments in accuracy across both reactive and proactive tasks^41^. Thus, current results preliminarily suggest that hypercapnia may potentially improve cognitive performance during tasks which require the parsing of conflicting information (i.e., stimulus-driven), but may not improve performance on tasks that require the planned/anticipatory allocation of cognitive resources^39, 44^. Importantly, the reduction in interference during reactive cognitive control requires replication in a controlled clinical design with placebo CO_2_ administration to rule-out practice effects, as well as testing in other cognitive domains such as vigilance and processing speed to investigate whether findings generalize to other domains.

Limitations of the current study include moderate sample sizes within each group that may have reduced power to detect more subtle differences in protein efflux and/or depletion as a function of TBI (small to medium effect sizes), as well as potential group differences in bulk CSF flow metrics (e.g., amplitude a statistical trend). Similarly, the current study followed recent recommendations to analyze TBI as a spectrum rather than based on traditional labels of “mild” or “moderate-to-severe” TBI^38, 63^. However, we were underpowered to examine for known imaging differences (e.g., greater atrophy and cerebrovascular impairment in more severe injuries) and possible differences in protein efflux based on injury severity, representing a critical future direction. As a result, means and standard deviations are provided in Supplemental Results for all proteomic measurements based on sub-group membership.

Second, the measurement of bulk CSF flow via BOLD imaging in humans is both a surrogate and indirect measurement of glymphatic functioning. Like most time-dependent imaging techniques, it is affected by multiple other physiological confounds (motion, respiratory patterns, etc.). However, multiple rodent studies suggest that bulk CSF flow represents a proxy measure for the penetration of CSF into parenchymal spaces^6–8^, suggesting that it may represent a reasonable surrogate of glymphatic function. Moreover, the methodologies used in the current study to manipulate cerebral blood flow (hypercapnia), proxies for CSF flow (T2* effects within the most inferior slice) as well as surrogates for waste clearance (commercially available proteomics) provide immediate clinical translation relative to the more complex methods/markers that have proven to be feasible only in animal models to date^4, 6, 7^. Finally, additional preclinical studies are required to determine if the decrease in neural abundant protein concentrations in the blood truly reflects a reduction in parenchymal protein levels, or is instead reflective of a reduced systemic clearance or temporary change in cerebrovascular tone post-hypercapnia.

In summary, there are currently no approved therapies for TBI outside of neurosurgery and acute symptom management. Current results suggest that CO_2_-induced LFHO are sufficient for increasing protein efflux, and may result in a brief period of parenchymal protein depletion before returning to pre-hypercapnia levels. The putative underlying mechanism of action for CO_2_ therapy (i.e., enhanced advective clearance through vascular-enhanced bulk CSF flow) harnesses normal brain physiology, and may therefore have widespread clinical applications for treating multiple neurodegenerative disease states relative to drugs that specifically target abnormal tau, amyloid or alpha-synuclein aggregation.

## Acknowledgements

We would like to acknowledge Haleemah Jackson for help with blood collection.

## Author Contributions

Conceptualization: ARM

Methodology: ARM, TVW, SGR, DSK, JML, KW, UN, VZ, RM

Investigation: ARM, TVW, SM, UN, JW, PC Visualization: ARM, TVW, SGR, UN Funding acquisition: ARM

Writing – original draft: ARM, TBM, RM Writing – review & editing: All authors

## Data Availability

Access to all data is available upon request to the corresponding author (ARM). Access to biospecimens from the study will be evaluated on a case-by-case basis, but are currently available.

