## Supplemental Materials for "Oscillating Hypercapnia Induces Neural Abundant Protein Efflux and Potential Depletion in Health and Chronic Traumatic Brain Injury"

***Supplemental Methods***

*Cognitive Control Task Description*

A multimodal task examined both reactive and proactive cognitive control pre- and post-hypercapnia, with previous studies demonstrating robust between condition differences in both healthy controls and participants with TBI[1, 2]. The cognitive control task started with a simultaneously presented multisensory (auditory and visual) cue (exemplary visual angle = 7.69°; duration = 300 ms) on either a black or a gray background. The cue prompted participants to attend to a specific sensory modality ("HEAR" = attend-auditory; "LOOK" = attend-visual), and was followed by simultaneously presented congruent or incongruent multisensory numeric stimuli ("one", "two" or "three"; duration = 300 ms) that served as the targets. The delay between the multisensory cue and the first target was at a fixed interval of 1000 ms, while the inter-stimulus interval between targets was fixed at 1200 ms (inter-target interval = 1500 ms).  To strengthen semantic relationships with auditory modality, visual stimuli were presented in word rather than numerical ("three" versus "3") form[3]. All multisensory stimuli (cues and targets) were presented foveally (eye-centered) and delivered binaurally through headphones (head-centered). To ensure spatial correspondence between auditory and visual stimuli, participants were required to maintain visual fixation on a centrally presented cross throughout all trial-types.

The background color upon which stimuli were presented further indicated trial type. A black background indicated that participants should press the button corresponding to the target presented in the attended modality (0-back trials) with their right hand while ignoring information in the unattended modality. This trial therefore measured reactive cognitive control and cognitive interference (i.e., large difference = poor reactive control) via subtraction of congruent from incongruent trials on the 0-back condition[1]. In contrast, a gray background signaled the performance of a 1-back task to measure proactive cognitive control. During gray trials, participants were required to hold the attended stimuli in working memory for 1.5 seconds (inter-target interval), ignore the unattended modality, and to only respond to the target upon the subsequent trial. Proactive cognitive control was quantified by subtracting congruent 0-back trials from congruent 1-back trials, with the expectation that more negative numbers indicate enhanced proactive cognitive control[4].

Stimuli were presented in blocks that pseudo-randomly varied between attend-auditory and attend-visual conditions, between the 0-back and 1-back conditions, and between congruent and incongruent multisensory stimulus conditions (8 trial types). In each block, 9 target stimuli were presented with a 1500 ms interval, corresponding to a frequency of 0.66 Hz. A total of 40 blocks were presented across 3 separate runs, with each run lasting for approximately 4.5 minutes. The inter-block intervals were jittered (2800, 3600, 4400 ms) to minimize temporal expectations.

*Imaging Acquisition and Processing*

All imaging data were acquired at a single site using a Siemens 3 Tesla PRISMA fit MRI scanner with a 32-channel head coil. High resolution $T_1$-weighted [repetition time (TR)=2530 ms; echo times (TE)=1.64, 3.52, 5.40, 7.28, 9.16 ms; inversion time (TI)=1200 ms; flip angle=7°; number of excitations (NEX)=1; slice thickness=1 mm; field of view (FOV)=256 mm; matrix size=256 × 256; isotropic voxels = 1.0 $mm^3$, 192 slices] scans were obtained and reviewed by a board-certified neuroradiologist blinded to diagnosis. Whole-brain BOLD data were obtained using a single-shot, gradient-echo echoplanar pulse sequence (575 volumes; runtime = 4 min 43 s; TR=480 ms; TE=29 ms; flip angle=44°; multiband acceleration factor=8; NEX=1; slice thickness=3 mm; field of view=248 mm; matrix=82×82; voxel size=3.0 $mm^3$; 56 slices). A single-band reference image was acquired to facilitate registration of BOLD data with the $T_1$–weighted image. Two spin-echo field mapping sequences with identical resolution and reversed phase encoding directions (A→P; P→A) were collected to mitigate susceptibility-related distortion.

*Physiological Modeling and Analysis*

MATLAB (MathWorks, Inc., Natick, MA; version R2014) was used to process raw $CO_2$ data similar to previous publications[5, 6], and included a 125 ms rolling maximum filter followed by post-collection recalibration (adjustment by two-point linear equation to expected ambient and gas mixture $CO_2$ floors). The majority of $ETCO_2$ peaks were identified algorithmically, with manual adjustment occurring as necessary. A spline function interpolated $ETCO_2$ data between peaks in conjunction with 10 s median filter smoothing, followed by resampling to match image acquisition parameters (repetition time). The resulting $ETCO_2$ regressor was Min-max normalized to approximate an amplitude of 1 across the entire timeseries, and shifted thirty-five times at 480 ms to create an array of regressors capable of modeling individual differences in physiology during the hypercapnia challenge.

*MR Data Analyses*

The first image of the BOLD timeseries were removed to account for $T_1$-equilibrium artifacts in addition to manufacturer's pre-programmed dummy scans. BOLD data were then processed using individual CSF and GM-specific pipelines (Supplemental Fig. S1) to determine vascular enhanced bulk CSF flow[7-9] and to characterize global GM cerebrovascular reactivity (CVR)[5, 6]. For the GM pipelines, anomalous timepoints were first identified and replaced using AFNI's despiking tool, followed by correction (temporal interpolation) for time-slice acquisition on a voxel-wise basis. Motion correction, performed in both 2D and 3D space, occurred next in relation to the single-band reference image. FSL's Topup algorithm was then used to correct for susceptibility induced field distortions. A global GM mask was next created from the $T_1$-weighted image (SPM12) using a 60% probability threshold, which was then down-sampled (3×3×3 mm) and spatially transformed to the single-band reference image (i.e., native space).

To determine CVR, the subject-specific GM mask was then multiplied by the preprocessed voxel-wise timeseries data to form a global GM timeseries vector. The array of participant-specific, time-lagged $ETCO_2$ regressors was then fit to the global GM vector. The unstandardized beta (i.e., amplitude) from regressor with the maximal fit (i.e., highest absolute Pearson correlation between $ETCO_2$ and global GM signal) was selected, followed by a Fisher's Z transformation of the Pearson correlation coefficient. The latency to maximal $ETCO_2$ fit was calculated by multiplying the index of maximal fit index by the associated lag time for the selected regressor. A constant of 7.2 seconds was added to CVR latency calculations to account for lung-brain[10, 11] and respirate-$CO_2$ recorder transit times (range 7.2-24.48 seconds).

To calculate vascular-enhanced bulk CSF flow resulting from $CO_2$-induced low frequency hemodynamic oscillations, the fully preprocessed voxel-wise data were simultaneously bandpass filtered (0.0008 to 0.10 Hz) and detrended using a $2^{nd}$ order polynomial function. The subject-specific GM mask was then multiplied by the band-pass filtered voxel-wise timeseries data to form a second global GM vector to characterize vascular-enhanced bulk CSF flow. The first 45 and last 15 images were truncated to account for edge artifacts associated with the bandpass filtering, followed by Min-max normalization of the gray matter signal. The band-passed GM data were then time-shifted at a resolution of repetition time 83 times to create an array of regressors (0 to 40.32 s). Numerical differentiation (N - (N-1)) was used to create an additional global GM derivative from the filtered data[7-9], which was then time-shifted 6 times to create a regressor array spanning from 0 to 3.36 s.

The minimal CSF-specific preprocessing stream included only time-slice correction, followed by band-pass filtering and detrending[7-9]. The most inferiorly collected BOLD slice was used to create a region of interest (ROI) that identified CSF within the cervical central canal inferior to the medulla oblongata. The ROI was generated via a semi-automated process, which was inspected by two independent raters, and manually edited when necessary. All raters achieved a minimum dice coefficient of 0.70 on a training dataset, and a third rater was involved for any significant inter-rater discrepancies. The subject-specific CSF ROI mask was then multiplied by the minimally preprocessed, band-passed filtered voxel-wise timeseries data to create a vector corresponding to changes in BOLD signal resulting from bulk CSF flow. From a physics perspective, CSF that has not been exposed to radiofrequency pulses has a higher T2* signal (i.e., flowing into the most inferior slice), whereas radiofrequency exposed CSF has a lower signal[12]. The first 45 and last 15 images of the bulk CSF flow vector were similarly truncated to reduce the impact of edge effects introduced by the bandpass filtering step.

To quantify vascular-enhanced bulk CSF flow, the array of participant-specific, time-lagged band-pass filtered global GM vectors (independent variables) were next fit to the bulk CSF flow vector using identical methods as the $ETCO_2$-global GM data. The latency for vascular-enhanced bulk CSF flow was calculated by multiplying the maximal fit index by the repetition time. The global GM derivative vector was also randomly shuffled 1000 times based on phase information (3dTsort in AFNI) and subsequently fit to the bulk CSF flow vector for each iteration. The correlation of the maximal fit between the global GM derivative and the CSF flow vector was assessed to see if it was greater than 97.5% of the random permutations (i.e., single tail from the null distribution) on a subject-specific basis as part of quality assurance steps.

The FreeSurfer (version 7.1.1) package was used to produce global tissue volume estimates from the $T_1$-weighted images[13]. These included total GM, WM and CSF volumes, which were subsequently normalized by total intracranial volume to correct for individual differences in head size. Brain age (secondary outcome) was also estimated using the $T_1$-weighted image and a deep learning network (DeepBrainNet) architecture[14] implemented in Python (v3.10) with AntsPy (v0.3.4). The algorithm was initially trained using a diverse cohort of 11,729 participants aged 3–95 years old from multiple imaging sites and thus is readily generalizable to multiple ages and disease types. Preprocessing steps for brain age estimation (Supplementary Fig. S2) included N4 bias field correction, brain extraction, and affine registration to the Montreal Neurological Institute (MNI) anatomical template. From the registered volume, 80 axial slices (range 45–125) were extracted and converted to 3-channel RGB format[15]. Brain age was predicted on a slice-by-slice basis using the trained model with TensorFlow and Keras (2.10). The predicted brain age difference (PAD) was calculated by subtracting actual subject age from the median value of all slice-wise predictions.

*Blood Processing*

Samples were centrifuged for 15 minutes at 1500 relative centrifugal force, aliquoted into 250 µl tubes, and immediately stored in an -80°C freezer. Protein quantifications were conducted blinded to group assignment.

#### Results

##### *Supplemental Results*

*Protein Efflux and Depletion*

Blood collection was successful in 216/220 (98.2%) attempts for individuals who completed the hypercapnia protocol (N=44). Eight of the collected samples were severely lysed, including 4 samples from a single HC (data not sent for processing). Blood samples were only successfully collected on 3/5 attempts for another HC. A total of 204 samples from 42 participants (20 HC, 9 mTBI, 13 msTBI) were therefore sent for protein quantification based on standard available well limits for duplicate processing on the Quanterix platform. A total of 1 baseline and 2 pre-hypercapnia samples were either not able to be collected or were lysed. Baseline and pre-hypercapnia samples were imputed for subsequent analyses based on age, sex and group membership given the low missingness and high fidelity of imputation models to explain variance (all models $R^2$>0.80). An LME model with plate as a nuisance covariate indicated trend Group differences at baseline for GFAP (F=3.32; *p*=0.077; Cohen's d=-0.60), NfL (F=3.71; *p*=0.062; Cohen's d =-0.63) and for brain derived tau (F=3.15; *p*=0.085; Cohen's d=-0.58) with non-significant effects for ptau217 (*p*>0.10; Cohen's d=-0.11).

*CVR and Bulk CSF Flow Coupling Results*

Extensive quality assurance protocols were conducted for the hypercapnia task on a run-by-run basis (N=44; 6 runs). The majority of $ETCO_2$ traces (250/264; 94.7%) exhibited clear evidence of at least 2 hypercapnic cycles per run. There was clear evidence of CVR within global GM (r>0.60) for 74.6% of runs, with an even higher number of runs (90.5%) exhibiting evidence of significant anti-correlation (>0.025% of null distribution) for $CO_2$-induced bulk CSF flow based on the GM derivative. Importantly, the majority of failed runs for both CVR (53/67) and $CO_2$-induced bulk CSF flow (15/25) occurred within the chronic TBI cohort, indicative of potential pathology across groups. CVR and $CO_2$-induced bulk CSF flow metrics (primary = fit and amplitude; secondary = latency) were summed across the 6 runs to facilitate group comparisons and analyses regarding biological mechanisms underlying protein efflux.

*Structural Imaging Results*

Blinded review of the $T_1$-weighted images indicated a significantly higher ($\chi^2$=6.844, $p$=0.009) percentage of abnormal scans for participants with TBI (8/22) relative to HC (1/22), predominantly as a result of encephalomalacia or presence of white matter hyperintensities.

### Supplemental Tables

Supplemental Table 1: Central tendency for blood protein data across sub-groups at each timepoint.

| Protein | Time | HC (N=20) | TBI (N=22) | mTBI (N=9) | msTBI N=13 |
|---|---|---|---|---|---|
| GFAP | B | 3.78±0.47 | 3.87±0.71 | 3.63±0.70 | 4.03±0.70 |
| | ΔPre | 0.04±0.10 | 0.05±0.11 | 0.11±0.08 | 0.02±0.12 |
| | Δ45 m | 0.18±0.17 | 0.18±0.20 | 0.19±0.20 | 0.17±0.21 |
| | Δ90 m | -0.04±0.24 | -0.01±0.18 | 0.05±0.14 | -0.03±0.19 |
| | Δ2.5 h | 0.05±0.27 | 0.07±0.19 | 0.11±0.23 | 0.04±0.16 |
| NfL | B | 1.72±0.62 | 1.95±1.19 | 1.64±0.84 | 2.17±1.36 |
| | ΔPre | -0.03±0.18 | 0.02±0.17 | 0.04±0.16 | 0.01±0.18 |
| | Δ45 m | 0.07±0.20 | 0.14±0.17 | 0.16±0.18 | 0.13±0.17 |
| | Δ90 m | -0.08±0.21 | -0.08±0.22 | -0.01±0.11 | -0.12±0.26 |
| | Δ2.5 h | -0.06±0.26 | -0.03±0.26 | 0.05±0.21 | -0.09±0.28 |
| BD tau | B | 1.74±0.44 | 1.77±0.43 | 1.85±0.55 | 1.71±0.33 |
| | ΔPre | -0.05±0.06 | -0.00±0.12 | 0.05±0.10 | -0.03±0.13 |
| | Δ45 m | 0.02±0.10 | 0.07±0.16 | 0.06±0.16 | 0.08±0.16 |
| | Δ90 m | -0.11±0.13 | -0.09±0.18 | -0.05±0.17 | -0.10±0.19 |
| | Δ2.5 h | -0.01±0.16 | -0.03±0.15 | -0.04±0.17 | -0.03±0.14 |
| ptau217 | B | -1.62±0.33 | -1.58±0.38 | -1.48±0.39 | -1.64±0.38 |
| | ΔPre | -0.05±0.10 | -0.07±0.16 | -0.12±0.20 | -0.03±0.12 |
| | Δ45 m | 0.02±0.13 | -0.06±0.25 | -0.13±0.18 | -0.02±0.28 |
| | Δ90 m | -0.08±0.17 | -0.16±0.16 | -0.17±0.20 | -0.15±0.13 |
| | Δ2.5 h | -0.04±0.13 | -0.17±0.19 | -0.21±0.19 | -0.13±0.19 |

Notes: Acronyms: HC=healthy control; TBI=traumatic brain injury; mTBI="mild" TBI; msTBI="moderate-to-severe" TBI; B=baseline; Δ=change score between the timepoint provided and baseline; GFAP=glial fibrillary acidic protein; NfL=neurofilament light chain; BD tau= brain-derived tau; ptau217= phosphorylated tau 217; All protein concentrations are presented as logarithm (log) of picogram per milliliter (pg/mL); All data are reported at mean plus standard deviation (denoted by ±). Baseline data are not presented as change scores.

#### Supplemental Figures

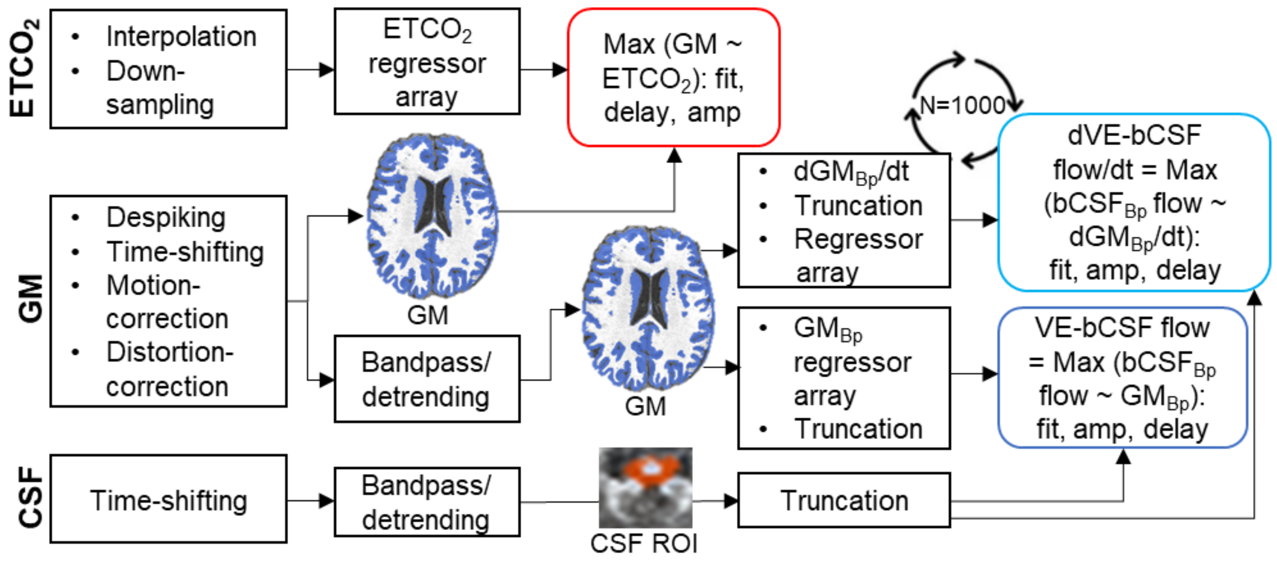


**Supplemental Fig. S1:** Schematic representation of the end-tidal carbon dioxide ($ETCO_2$), gray matter (GM) and bulk cerebral spinal fluid (CSF) flow processing pipelines. Per published recommendations, CSF data were processed with a minimal pre-processing pipeline that included a bandpass filter (Bp), detrending and truncation to remove edge artifacts. Gray matter data were more extensively pre-processed, and either excluded (cerebrovascular reactivity pipeline) or included bandpass filtering, detrending and truncation. Cerebrovascular reactivity was calculated by finding the maximum correlation between an array of time-shifted $ETCO_2$ regressors with the unfiltered gray matter data (red box), and included the associated fit (Pearson r), amplitude (amp) and delay parameters. Vascular-enhanced bulk CSF (VE-bCSF; blue box) flow was calculated by finding the maximum correlation between an array of time-shifted, bandpass filtered gray matter regressors with the filtered bulk CSF flow (bCSF flow) data , as well between time-shifted gray matter derivative data ($dGM_{Bp}/dt$; light blue box) with the filtered bulk CSF flow data. The derivative pipeline included randomly shuffling the timeseries data and permuting the regression

1000 times to develop a distribution from which to conduct quality assurance assessments on a subject-by-subject basis.

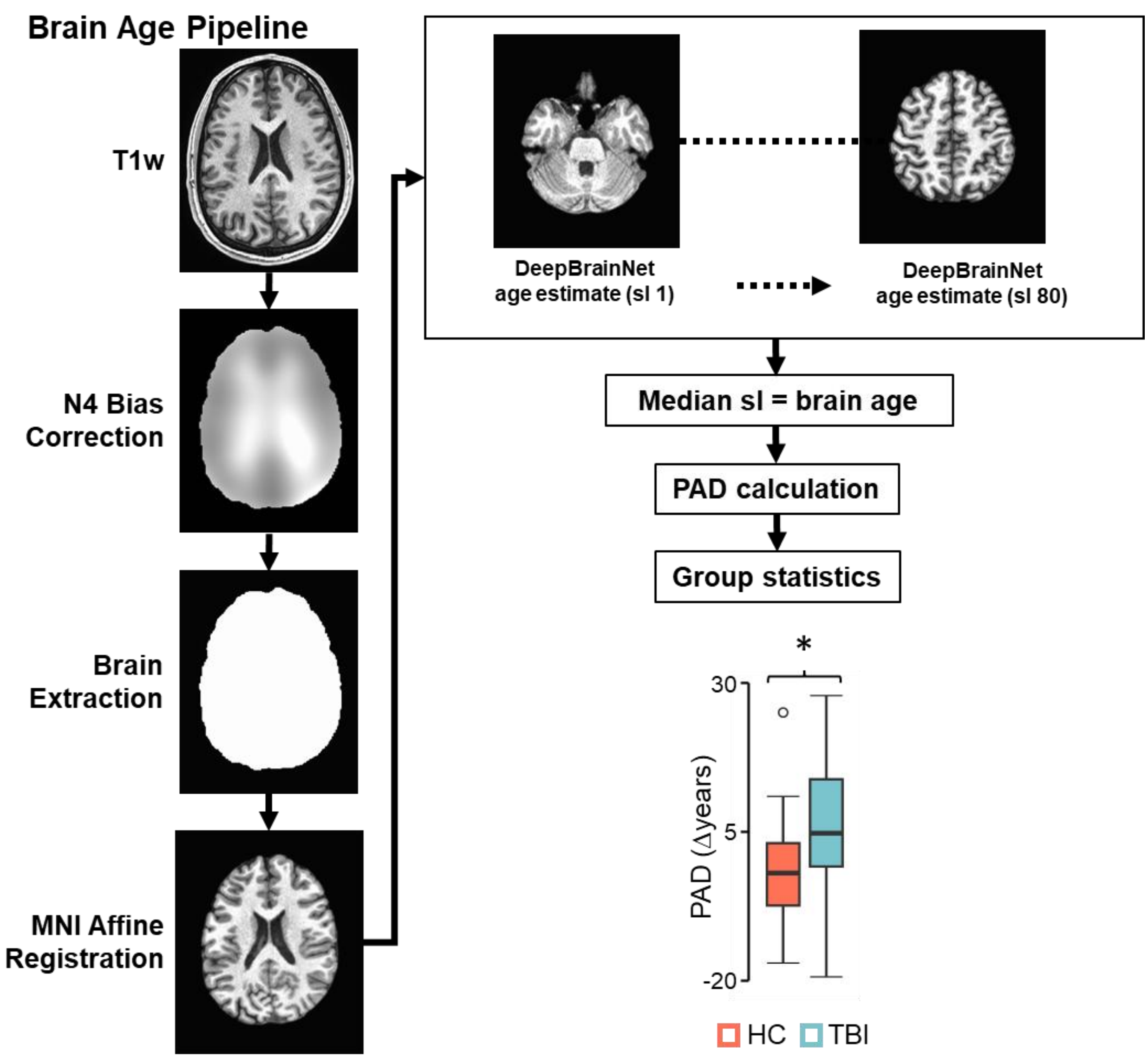


**<u>Supplemental Fig. S2:</u> Brain age methods and results.** Raw $T_1$-weighted (T1w) images were normalized to Montreal Neurological Institute (MNI) space, and brain age was estimated from the median of 80 slices (sl). Participants with chronic traumatic brain injury (TBI: light blue) exhibited significantly (group effects denoted by asterisk) increased predicted brain age difference (PAD) relative to matched healthy controls (HC: coral).

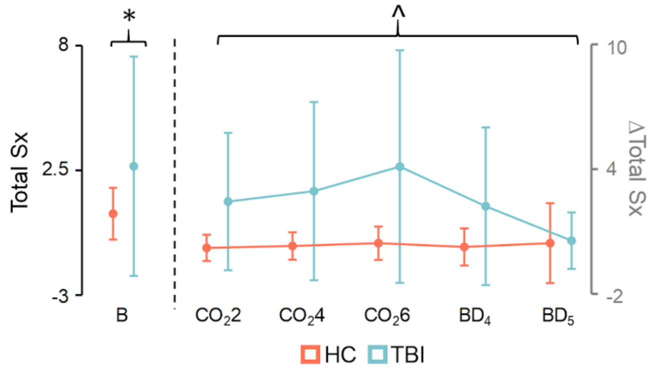


**<u>Supplemental Fig. S3:</u>** A depiction of total symptom (Sx) burden for chronic traumatic brain injury (TBI; light blue) participants and healthy controls (HC; coral) for the baseline (B) or the change between baseline and each of the subsequent five symptom assessments ($CO_2$=hypercapnia challenge followed by magnetic resonance imaging run number; BD=blood draw). Total symptom burden was significantly greater for chronic TBI participants at baseline (denoted by asterisk), and symptoms peaked across both groups following the last hypercapnia challenge (denoted by carrot symbol) relative to other assessments prior to quickly returning to baseline levels (error bars = standard deviation).
